# Dramatic expansion of microbial groups that shape the global sulfur cycle

**DOI:** 10.1101/166447

**Authors:** Karthik Anantharaman, Sean P. Jungbluth, Rose S. Kantor, Adi Lavy, Lesley A. Warren, Michael S. Rappé, Brian C. Thomas, Jillian F. Banfield

## Abstract

The biogeochemical cycle of sulfur on Earth is driven by microbial sulfate reduction, yet organisms from relatively few lineages have been implicated in this process. Recent studies using functional marker genes have detected abundant, novel dissimilatory sulfite reductases that confer the capacity for microbial sulfate reduction and could do not be affiliated with known organisms. Thus, the identity of a significant fraction of sulfate reducing microbes has remained elusive. Here we report the discovery of the capacity for sulfate reduction in the genomes of organisms from twelve bacterial and archaeal phyla, thereby doubling the number of microbial phyla associated with this process. Eight of the twelve newly identified groups are candidate phyla that lack isolated representatives, a finding only possible given genomes from metagenomes. Two candidate phyla, *Candidatus* Rokubacteria and *Candidatus* Hydrothermarchaeota contain the earliest evolved genes. The capacity for sulfate reduction has been laterally transferred in multiple events within some phyla, and a key gene potentially capable of switching sulfur oxidation to sulfate reduction in associated cells has been acquired by putatively symbiotic bacteria. We conclude that functional predictions based on phylogeny will significantly underestimate the extent of sulfate reduction across Earth’s ecosystems. Understanding the prevalence of this capacity is integral to interpreting the carbon cycle because sulfate reduction is often coupled to turnover of buried organic carbon. Our findings expand the diversity of microbial groups associated with sulfur transformations in the environment and motivate revision of biogeochemical process models based on microbial community composition.

---

The cycling of sulfur is one of Earth’s major biogeochemical processes. Sulfate reduction may be an early evolved microbial metabolism, given evidence for biological fractionation of sulfur isotopes around 3.5 billion years ago^1^, and it remains an important metabolic platform for anaerobic life^2^. In natural ecosystems, human microbiomes and engineered systems, this process is important because the product hydrogen sulfide (H_2_S) is toxic^3^, can corrode steel^4^, and sour oil reservoirs^5^. Overall, sulfate reduction is a primary driver in the carbon cycle, and is responsible for conversion of ~30% of the organic carbon flux to CO_2_ in sedimentary environments^6^. Importantly, the coupling of sulfate reduction to oxidation of H_2_, small chain fatty acids or other carbon compounds limits the availability of these substrates to other organisms and alters the energetics via syntrophic interactions^7^. All of these processes also impact methane production. Given the many reasons why the biological conversion of sulfate/sulfite to sulfide is important, it is vital that we understand which organisms can carry out the reactions and the pathways involved.

Dissimilatory sulfite reductase (dsr) genes confer bacteria and archaea the ability to grow via reduction of sulfite and can function in reverse in some organisms that oxidize sulfur^8,9^. The phylogenetic distribution of organisms with dsr genes has been considered to be quite limited^10^. The recent availability of thousands of genomes from organisms belonging to many newly sampled phyla has provided the opportunity to test for the presence of dsr genes in bacteria and archaea that have not previously been associated with dissimilatory sulfur metabolism^11^.

## Results and Discussion

We analyzed genomes reconstructed from metagenomic sequence datasets recovered from five distinct terrestrial and marine subsurface environments. The sampling sites included an aquifer adjacent to the Colorado River, USA^12^, a deep subsurface CO_2_ geyser in Utah, USA^13^, a deep borehole in Japan^14^, an acidic sulfide mine waste rock site in Canada^15^, and deep subseafloor basaltic crustal fluids of the hydrothermally active Juan de Fuca ridge flank in the Pacific Ocean^16^. We identified dsr genes in 122 near-complete microbial genomes (**Supplementary Table 1**). Phylogenetic analyses using a set of 16 concatenated ribosomal proteins (RP) and the small subunit ribosomal (SSU) RNA gene show that these genomes belong to organisms from 16 distinct phylum-level lineages (**Table 1**), 12 of which were not known to have dsr genes^10^. In addition, we identified anaerobic sulfite reductase (asr) genes required for sulfite reduction in two bacterial groups not previously reported to have this capacity^17^. All of the identified catalytic proteins (DsrA, DsrB, and AsrC) contained all conserved sulfite reductase residues and secondary structure elements for the formation of α helices and β sheets^18^ (**Supplementary Fig. 1, Supplementary Fig. 2, Supplementary Fig. 3**).

**Table 1.**
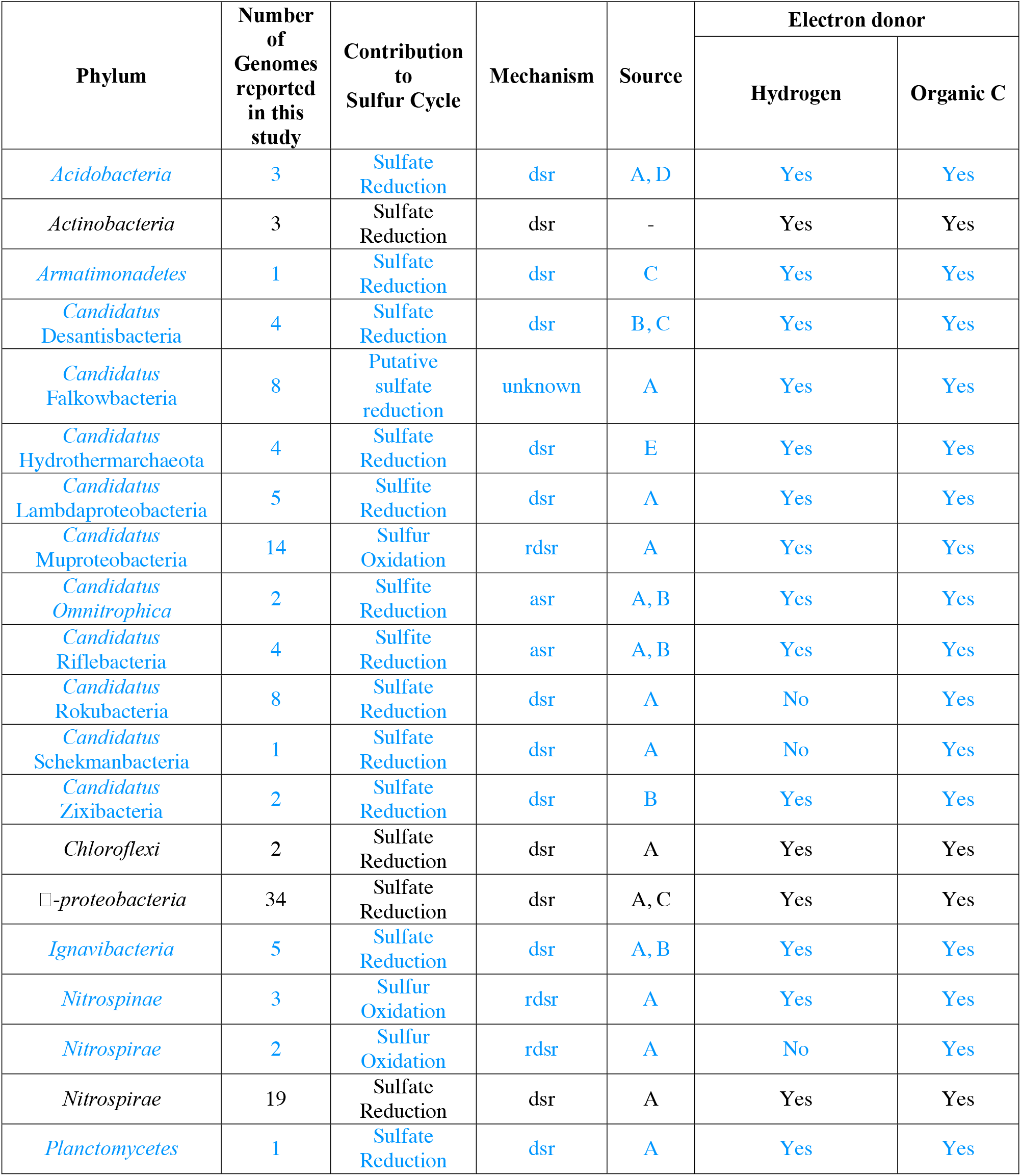
Details of lineages involved in dissimilatory sulfur cycling.

Given our interest in identifying organisms with the capacity to produce sulfide, we searched genomes for operons that contained genes encoding DsrD. This gene is considered a marker for sulfate reduction because it is absent in bacteria that use the reverse dissimilatory sulfate reduction (reverse-dsr) pathway for sulfur oxidation^19^. Although the exact function of the DsrD protein is unclear, the presence of winged-helix domains in its structure and its association with other core proteins of the dsr complex (*dsrABC*) suggest a regulatory role in bacterial sulfate reduction^20^. We identified 78 genomes that encode at least *dsrABCD* (**Supplementary Fig. 4**). A multiple alignment of DsrD sequences confirmed highly conserved residues, indicating that the proteins are likely active (**Supplementary Fig. 5**). These putative sulfate/sulfite reducing microorganisms affiliate with eight distinct phyla not previously reported to be capable of these processes. Four are phyla with isolated representatives (*Acidobacteria*, *Armatimonadetes*, *Ignavibacteria*, *Planctomycetes*) and four are considered candidate phyla due to the absence of isolated representatives (*Candidatus* Zixibacteria, *Candidatus* Schekmanbacteria, *Candidatus* Desantisbacteria, *Candidatus* Lambdaproteobacteria) (**Fig. 1**). The asr pathway for sulfite reduction was found in members of two candidate phyla, *Candidatus* Omnitrophica and *Candidatus* Riflebacteria (**Supplementary Fig. 6**).

**Fig.1.**
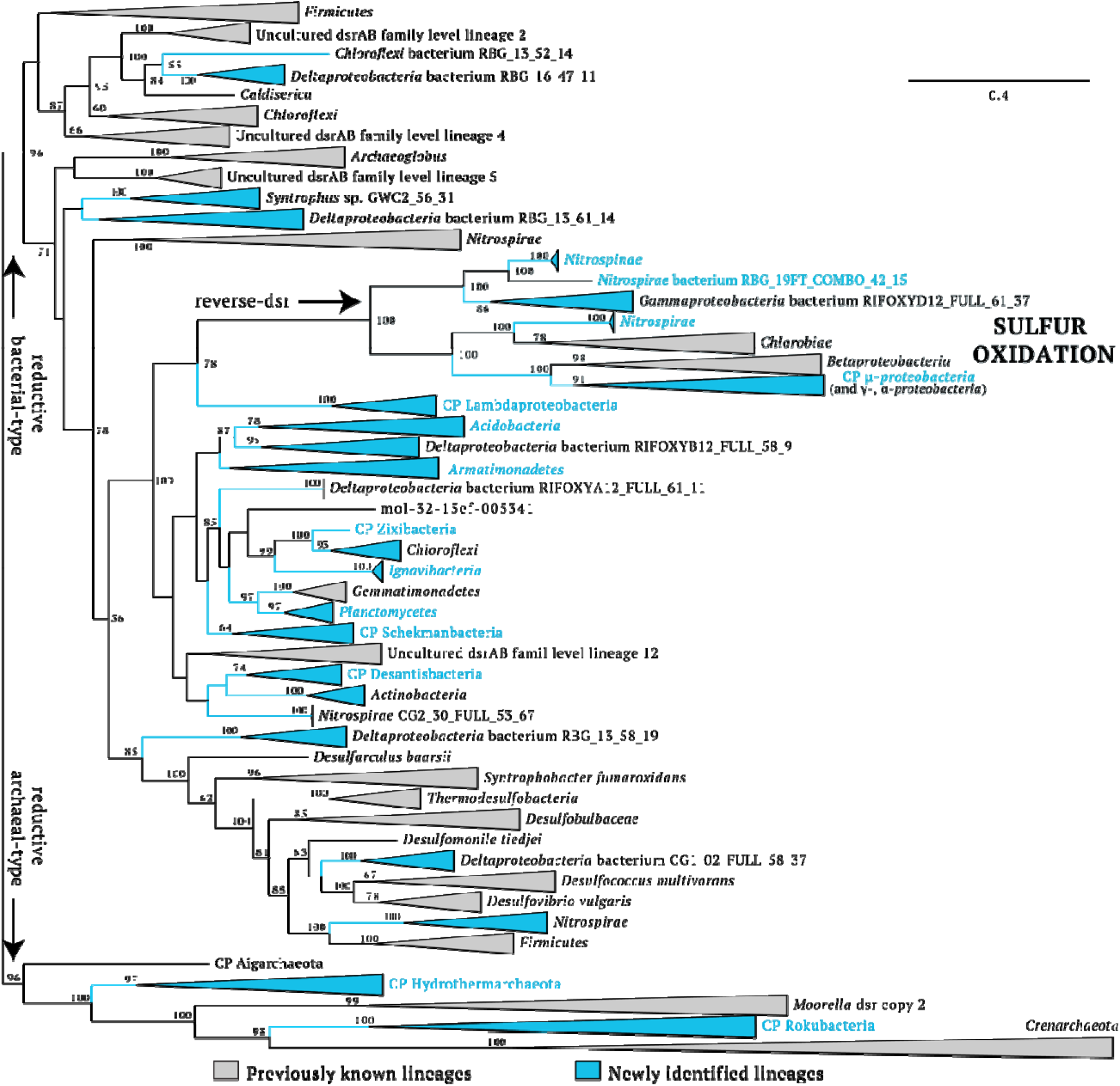
Phylogenetic tree of concatenated dsrAB proteins showing the diversity of organisms involved in dissimilatory sulfur cycling. Lineages in blue contain genomes reported in this study. Phylum-level lineages with first report of evidence for sulfur cycling are indicated by blue letters. Only bootstrap values >50 are shown. The complete tree is available with full bootstrap support values as Additional Data File S1.

Surprisingly, we identified *dsrD* genes in eight genomes of organisms affiliating with *Candidatus* Falkowbacteria, putatively symbiotic bacteria within the Parcubacteria superphylum of the candidate phyla radiation (CPR)^21^. There is no indication of the presence of other dsr genes in these genomes. Given the predicted close physical and metabolic interactions between CPR and their hosts, we suggest that this small protein could augment host metabolism, as sometimes occurs with viruses/phage and their hosts^22^. CPR are common in aquifers where conditions oscillate between oxic and anoxic^12^. Potentially, populations with the *dsrD* gene could maintain host function by enabling switching between sulfur oxidation to sulfate reduction as conditions change. The predicted Falkowbacteria DsrD protein sequences cluster with sequences from well characterized *Deltaproteobacteria* capable of sulfate reduction, suggesting that these CPR may have acquired the gene by lateral gene transfer (LGT) from this group (**Supplementary Fig. 7**).

Prior analyses have suggested the LGT events involving the catalytic dsr subunits A and B genes have occurred, but infrequently^9,10,23^. We used a concatenated *dsrAB* protein tree to reevaluate the extent to which LGT has influenced the organismal distribution of these genes (**Fig. 1**). We found that organism phylogeny is not a reliable predictor of the grouping of these sequences. In fact, phylogenetic evidence suggests that LGT events have introduced these genes into some phyla in multiple independent events (e.g., Nitrospirae sequences place in five distinct locations on the tree). However, almost all organisms lacking *dsrD* genes cluster together with organisms known to be sulfur oxidizers in the *dsrAB* tree. Based on this clustering, the group implicated in elemental sulfur oxidation now includes bacteria from three phyla: *Nitrospirae*, *Nitrospinae*, and *Candidatus* Muproteobacteria (**Fig. 1, Fig. 2**). Importantly, organisms from two candidate phyla, *Candidatus* Rokubacteria and *Candidatus* Hydrothermarchaeota lack *dsrD* genes but their *dsrAB* sequences cluster with ‘reductive archaeal-type’ dsr sequences found in thermophilic sulfite/thiosulfate-reducing *Crenarchaeota* and *Aigarchaeota*. The branch that includes sequences from *Candidatus* Rokubacteria, *Candidatus* Hydrothermarchaeota, *Crenarchaeota* and *Aigarchaeota* is basal within the *dsrAB* tree (**Fig. 1**). The phylogenetic placements of *Candidatus* Rokubacteria and *Candidatus* Hydrothermarchaeota with sulfite/thiosulfate-reducing archaea implicates these organisms in either sulfate or sulfite reduction.

**Fig.2.**
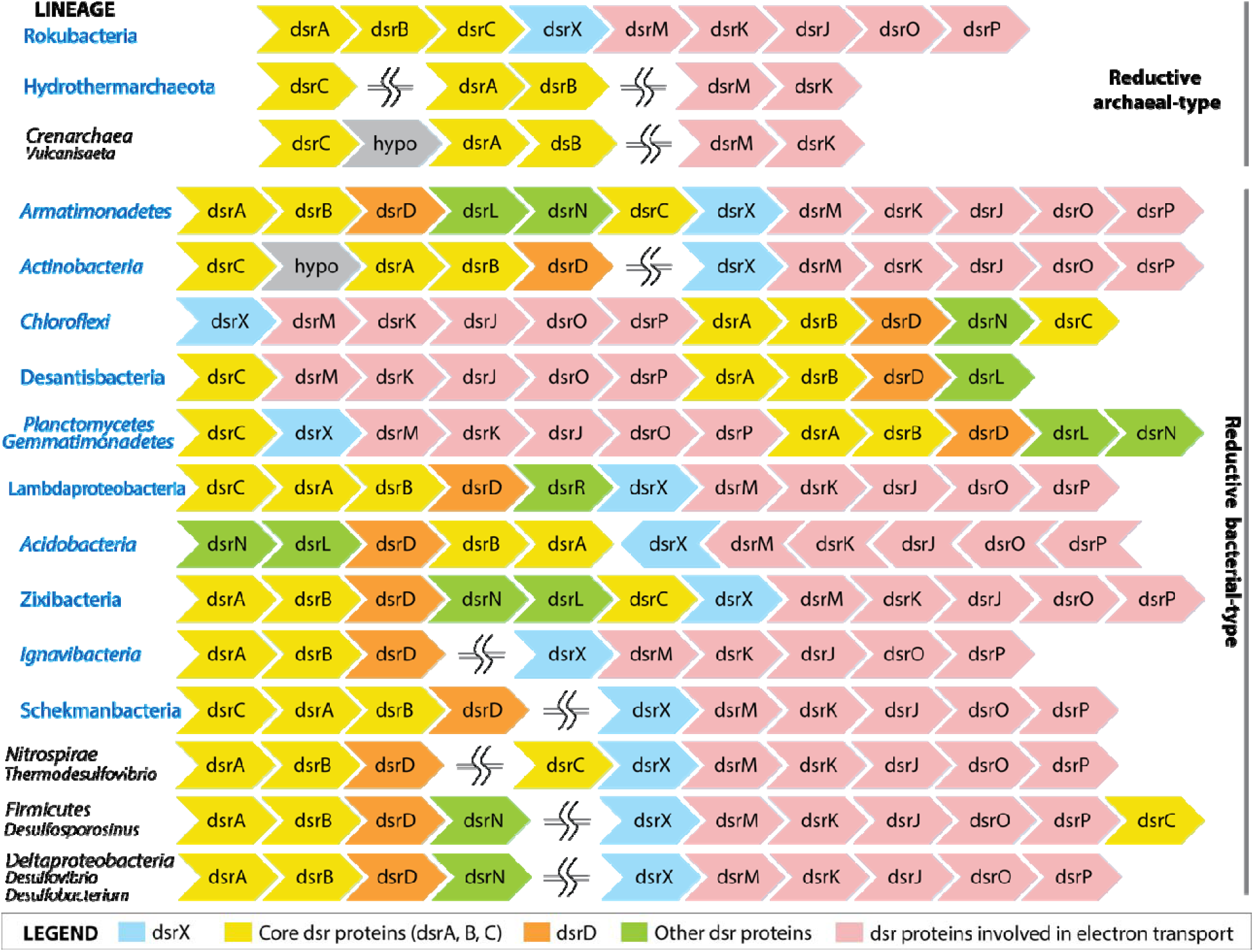
Dsr operon structure in previously reported (black names) and newly reported groups (blue names). Interestingly, and in contrast to the previously studied organisms for which the operon is interrupted (=SS=), the entire dsr pathway (including electron transport chain and ancillary proteins) is often encoded in a single genomic region.

To determine whether the organisms reduce sulfate vs. sulfite to sulfide we looked for the genes involved in the reduction of sulfate to sulfite, specifically adenosine phosphosulfate reductase subunits A, B (*aprAB*), sulfate adenylyl transferase (*sat*), and quinone-interacting membrane-bound oxidoreductase subunits A, B, C (*qmoABC*)^24–26^. Unlike *Crenarchaeota* and *Aigarchaeota* that can only reduce sulfite, *Candidatus* Rokubacteria have apr, sat, and the qmo genes that are required to gain energy from reduction of sulfate to sulfite. Surprisingly, phylogenetic analyses show that the *Candidatus* Rokubacteria and *Candidatus* Hydrothermarchaeota qmo (and *apr* and *sat*) genes cluster with sequences from other sulfate-reducing bacteria (**Supplementary Fig. 8, Supplementary Fig. 9, Supplementary Fig. 10**). Thus, we suggest that *Candidatus* Rokubacteria have a system for sulfate reduction to sulfide of hybrid origin, with ancient DsrA and B genes related to those found in archaea, and other components similar to bacterial sequences (**Fig. 3**).

**Fig. 3.**
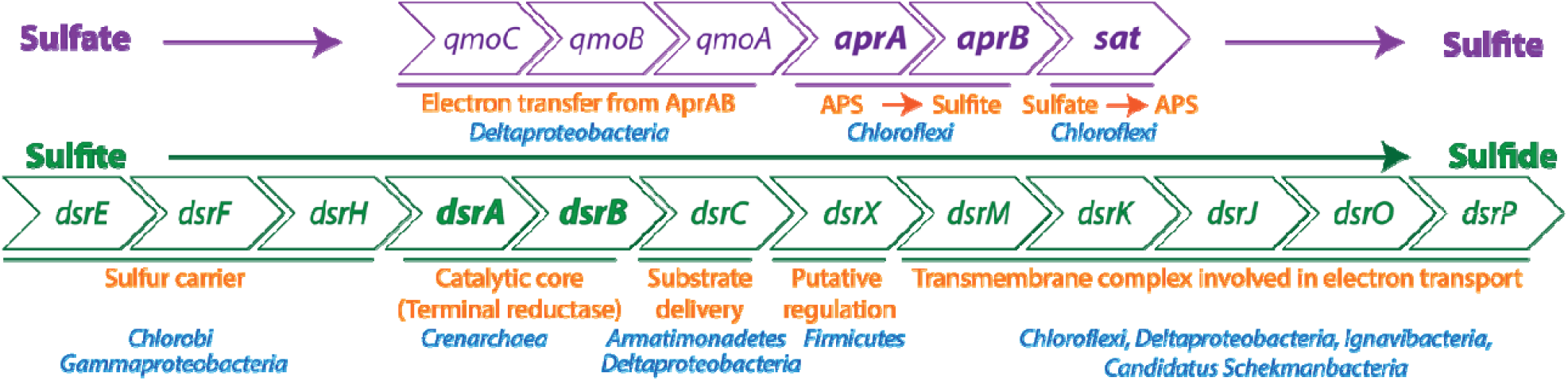
Dsr operon structure and enzymatic roles of proteins involved in sulfate reduction in *Candidatus* Rokubacteria. Purple: genes involved in sulfate reduction to sulfite. Orange: putative enzymatic roles of genes, blue: microbial lineages with closest homologs as determined by phylogeny/blast against NCBI Genbank. APS refers to adenosine-5’′-phosphosulfate. Green: genes involved in sulfite reduction to sulfide. This is the first case in which *dsrE*, *dsrF* and *dsrH* genes are present in organisms other than sulfur-oxidizing bacteria.

Given the lack of *dsrD* in *Candidatus* Rokubacteria and *Candidatus* Hydrothermarchaeota, we sought evidence for hypothetical genes in proximity to dsr genes that may be markers for the sulfate/sulfite reduction pathway. We identified a hypothetical gene that encodes for the N-terminal domain of an anti-sigma factor antagonist protein^27^ that almost always occurs within the operon encoding dsr genes (**Fig. 2**). This hypothetical protein is part of a protein family that includes the *Bacillus subtitlis* RsbT co-antagonist protein rsbRD, which are important components of the stressosome and function as negative regulators of the general stress transcription factor sigma-B^28^. This gene is unique to sulfide-producing organisms and is absent in sulfur-oxidizing organisms (except for the *Chlorobiae* clade) (**Supplementary Fig. 11**). The gene always precedes the electron transport components encoded by *dsrMKJOP* genes and is fused with *dsrM* in some organisms (**Supplementary Fig. 12**). From structural predictions and conserved motifs, we hypothesize that it likely performs a regulatory function (**Supplementary Fig. 13**). We refer to this newly identified gene as ‘dsrX’ and suggest that it may serve as an additional marker for genome-based prediction of sulfate or sulfite reducing metabolism.

In order to understand the energy metabolism and ecology of these novel sulfate reducing organisms, we investigated potential electron donors for sulfate/sulfite reduction. Specifically, we targeted genes involved in the oxidation of hydrogen^29^ (Ni-Fe hydrogenase groups I, IIa, IIb, IIIa, IIIb, IIIc, IIId) and transformation of organic carbon compounds (Genes involved in breakdown of Cellulose, Hemicellulose, Chitin, Pectin, Starch, Amino sugars, Other monosaccharides and polysaccharides)^30^. Our analyses show that organisms from 10 of the 12 sulfate reducing lineages possess the ability to utilize hydrogen as an electron donor for sulfate reduction (**Supplementary Table 2**). On the other hand, organisms from all 12 lineages possessed the ability to breakdown organic compounds although the diversity of genes encoding for specific carbohydrate active enzymes varied greatly across phyla (**Supplementary Table 3**). We propose that sulfate reduction by organisms from these newly identified lineages likely serves an important control on the transformation of organic carbon in the terrestrial and marine subsurface.

By the Proterozoic Eon, sulfate reduction had become a significant biological process in the oceans^31,32^. Based on phylogenomic arguments and isotopic records, it was suggested that the capacity to reduce sulfite to sulfide emerged in thermophilic archaea around 3.5 billion years ago, and that mesophilic sulfate reducers evolved only after the rise in atmospheric oxygen level^1,33^. Our findings indicate a complex evolutionary history of this capacity involving extensive LGT of dsr genes. Consequently, it may be impossible to constrain the lineage in which this metabolism first appeared. The ability to reduce sulfate/sulfite is now predicted in a much wider diversity of mesophilic bacterial and archaeal groups than was recognized previously. We conclude that many groups of microorganisms now known to have genes involved in dissimilatory sulfur metabolism impact biogeochemical processes in marine and terrestrial sediments, aquifers, wetlands, methane seeps, coastal marshes and estuaries, as well as agricultural and human microbiomes. Many are organisms from well-studied phyla, but still novel at the Genus to Class levels, but others are organisms from candidate phyla known only based on their genomes. The results underline the value of genomic analyses for prediction of key ecosystem capacities that cannot be made based on rRNA gene surveys and motivate targeted cultivation strategies for organisms currently lacking laboratory tractable representatives.

## Methods

### Sample Collection and Data Processing

Details of sample collection (Sampling, DNA Extraction), individual sample geochemical measurements, and data processing (DNA sequencing, Assembly, Annotation, Binning, Genome Completion Estimates) are described in detail elsewhere^12–16^.

### Identification of sulfate reducing organisms

Genome-specific metabolic potential for sulfate reduction was determined in an iterative manner by (A) Searching all predicted ORFs in a genome with hmm profiles for *dsrA* and *dsrB* from TIGRfam^34^, and *dsrD* from Pfam^35^ using hmmscan v3.1b2^36^, and (B) Generation of custom hmm profiles for *dsrA*, *dsrB* and *dsrD* using hits generated from step (A) and searching all predicted ORFs again for the above genes. For generation of custom HMM profiles, reference sequences and identified genes from step (A) were aligned using MUSCLE v3.8.31^37^ with default parameters followed by manually trimming the start and ends of the alignment. The alignment was converted into Stockholm format and databases were built using hmmscan^36^. Individual noise and trusted cutoffs for all HMMs were determined by manual inspection and are built into the custom HMM profiles.

### Sequence alignment and phylogeny

Phylogenetic analyses were performed as follows:

Each individual gene (*dsrA*, *dsrB*, *dsrC*, *dsrD*, *aprA*, *aprB*, *dsrX*, *qmoA*, *qmoB*, *sat*) was aligned along with reference sequences using MUSCLE^37^ with default parameters. All alignments were manually refined by trimming the start and ends and removing all columns with >95% gaps. For generation of concatenated alignments (*dsrAB*, *qmoAB*, and *aprAB*), individual alignments were concatenated in Geneious version 7^38^. In construction of the concatenated qmo tree, only subunits A and B were used since subunit C is not universally present in sulfate reducing organisms, being absent in sulfate reducing archaea. All phylogenetic analyses were inferred by RAxML v8.0.26^39^ implemented by the CIPRES Science Gateway^40^. RAxML was called as follows:

For *asrABC*, *dsrD*, *sat*, *dsrX*, *qmoAB* trees:

raxmlHPC-PTHREADS -s input -N 1000 -n result -f a -p 12345 -x 12345 -m PROTGAMMAGTR.

For *dsrAB*, *aprAB* trees:

raxmlHPC-HYBRID -s input -N autoMRE -n result -f a -p 12345 -x 12345 -m PROTGAMMAGTR.

### Conserved residues and motifs

Conserved residues and motifs in DsrA, DsrB, DsrC, AsrC, and DsrD proteins were identified by aligning the identified genes from all 122 genomes in this study with reference proteins^18,20,41^. Specific residues highlighted in **Supplementary Fig. 1**, **Supplementary Fig. 2**, **Supplementary Fig. 3**, **Supplementary Fig. 5**, and **Supplementary Fig. 14** were identified in *Desulfovibrio vulgaris*^42^.

### Structural models

We selected the DsrX proteins identified in *Desulfovibrio vulgaris* (WP_012611240) and Candidatus Rokubacteria CSP1-6 (KRT71371) for structural modeling. Protein models were predicted using the I-TASSER suite^43^. The models shown in **Supplementary Fig. 12** are the top predicted models out of the top five I-TASSER simulations. Both DsrX proteins used the identical top threading template from the sporulation inhibitor protein pXO1-118 from *Bacillus anthracis*^44^.

### Analyses of electron donors for sulfate reduction

Analyses of putative electron donors were centered around hydrogen and organic carbon compounds (carbohydrates). For identification of the potential for hydrogen oxidation, hmm searches were conducted by searching all predicted ORFs against individual HMM profiles for nickel-iron hydrogenases from Groups I, IIa, IIb, IIIa, IIIb, IIIc, and IIId. All hits above the noise cutoffs were inspected manually.

For identification of carbohydrate substrates for sulfate reduction, all predicted ORFs were searched against the CAZY HMM database^30^. Pre-filtering of hits was conducted using the following cutoffs: coverage: 0.40; e-value: 1e-18. To determine the specificity of enzymes, we established a set of 84 distinct reactions involving 189 enzyme families that allowed us to track specific substrates and products. All hits to Glycosyltransferases (GT) and Carbohydrate Binding Modules (CBM) were excluded from this analysis due to high incidence of false positives and/or difficulty in determining substrate specificity.

### Data availability

NCBI Genbank, BioProject, BioSample, and Taxonomy ID (TaxID) accession numbers for individual genomes are listed in Supplemenatry Table 1. Genomes are also available through ggKbase: http://ggkbase.berkeley.edu/novel_sulfate_reducers(ggKbase is a ‘live’ site, genomes may be updated after publication). The JdFR-17, JdFR-18, and JdFR-19 genomes are also available through the Integrated Microbial Genomes and Microbiomes database (IMG) through Genome IDs: 2728369317, 2728369320, 2728369322. Hmm databases used in this study are available from https://github.com/banfieldlab/metabolic-hmms. Additionally, ARB databases for *dsrA*, *dsrB*, and concatenated *dsrAB* genes are available from https://github.com/banfieldlab/dsrAB-ARB-db. The authors declare that all other data supporting the findings of this study are available within the article and its supplementary information files, or from the corresponding author on request.

## Acknowledgements

We thank David Burstein for inputs into hmm analysis and Christopher T. Brown for helpful discussion. This work was supported by Lawrence Berkeley National Laboratory’s Sustainable Systems Scientific Focus Area funded by the U.S. Department of Energy, Office of Science, Office of Biological and Environmental Research under contract DE-AC02-05CH11231.

## Additional information

**Supplementary information is available for this paper**

**Correspondence and request for materials** should be addressed to K.A.

